# Unraveling the Folding Dynamics of DNA Origami Structures

**DOI:** 10.1101/2025.07.11.664431

**Authors:** Meysam Mohammadi Zerankeshi, James Houston, Ogochukwu K.U. Elisha-Wigwe, Abi Sachi, Alexander E. Marras

## Abstract

Achieving high folding yield remains a major challenge in DNA origami, particularly as structures increase in complexity and scale. Here, we investigate how DNA origami design influences folding yield and kinetics using a combination of real-time fluorometry, gel electrophoresis, electron microscopy, and theoretical analysis. Results reveal a balance of the free energy changes from loop formation and hybridization that govern nucleation of nanostructure assembly, while the extent of cooperativity determines the overall assembly behavior. We measure the effect of structural complexity, staple design, and scaffold design on each energetic parameter, folding yield, and kinetics. We show that the scaffold crossover pattern determines the extent of cooperativity and subsequent folding kinetics, where fewer scaffold crossovers result in more cooperative folding. We also demonstrate that limiting the number of crossovers per staple should be prioritized over extending staple binding domains. The entropic penalty dominates the lower energy binding, disrupting folding. Finally, we demonstrate a 1-2 hour focused annealing ramp strategy that can increase yield up to 17% relative to traditional multi-day ramps. Optimizing energy changes and the contribution of cooperativity through design can significantly enhance folding yield and assembly time, particularly for complex structures, aiding the design and assembly of large-scale materials.

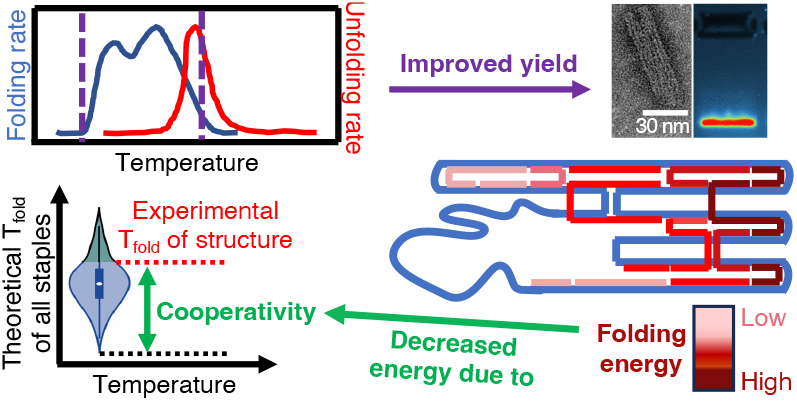

---

The development of DNA-based nanomaterials through the controlled self-assembly of engineered nucleotide sequences is responsible for the best geometric precision of any soft material. Recent advances using DNA origami include shells for trapping viruses,^1^ a ratchet-driven molecular rotor,^2^ and a tunable molecular force sensor.^3^ Further assembly efforts expand DNA origami technology to larger scales through hierarchical assembly of nanostructures,^4–7^ requiring high yield and larger scale production of nanostructures. However, producing DNA nanostructures rapidly and at high yields are still significant challenges that should be addressed to reach the full potential of DNA origami.^8–10^ A comprehensive understanding of the complex DNA origami assembly process will bring us closer to this goal.

DNA origami is fabricated via a long single-stranded DNA ‘scaffold’ that is folded into predetermined 2D or 3D shapes by many short ‘staple’ DNA oligonucleotides that are the complementary counterparts of the scaffold sequence.^11,12^ The result can be a vast range of nanoscale devices and structures with applications spanning drug carriers,^13,14^ mechanically functional nanomaterials,^15–17^ biosensor platforms ^18–20^ for arrangement of molecules,^21,22^ and much more. Unfortunately, with complex designs some applications become limited due to poor fabrication yield and the typical requirement of a multi-day fabrication process, but recent studies are revealing paths to overcome these barriers. Sobczak *et al.*^8^ compared the folding behavior of DNA origami objects to reveal the significant role of staple routing and cooperativity. By employing constant temperature annealing, they enhanced the folding yield and sped up the fabrication time. Martin and Dietz^23^ showed that emphasizing longer continuous staple domains improved folding yield. Recently Aksel *et al.*^9^ introduced a computational framework to optimize staple routing based on robust thermodynamic models to increase yield and precision of DNA origami fabrication. In this study, machine learning enabled auto-breaking of staples of any given design that would result in the highest yield. Furthermore, DeLuca *et al.*^24^ employed a mesoscopic simulation model that revealed a multi-step folding process for DNA origami, where structures initially zip into a partially folded precursor before crystallizing into their final form. In another study,^25^ Monte Carlo simulations showed that nucleation barriers of staple binding are design-dependent, with coaxial stacking of adjacent staples playing a critical role in assembly time and yield. Even with advancements in understanding the DNA origami folding process, the intricate relationship between nanostructure design and assembly behavior for complex structures remains vague. Many parameters including scaffold routing, staple break strategies, and folding protocols influence assembly through changes in local and global free energy and the role of cooperativity. Further focused studies are required to fully characterize this process.

There are three primary thermodynamic factors that regulate the binding of a staple to the associated section of the scaffold, and hence, the total assembly behavior of the nanostructure,^26–28^ as recently described in Aksel *et al.*^9^ and depicted in Figure 1a. ΔG_hyb_ is the free energy change due to scaffold-staple hybridization, which is a favorable term because the formed duplex has lower energy than the single-stranded counterparts. The second term, ΔG_loop_, is the energy cost to the system from decreasing entropy by folding the scaffold and overcoming electrostatic repulsion, an unfavorable energy change. The third term is the energy required for the diffusion of the corresponding staple, shown by ΔG_diffusion_. These energy changes collectively contribute to the total free energy change (ΔG_total_) of the staple during binding, playing a crucial role in the assembly process.^29,30^

**Figure 1.**
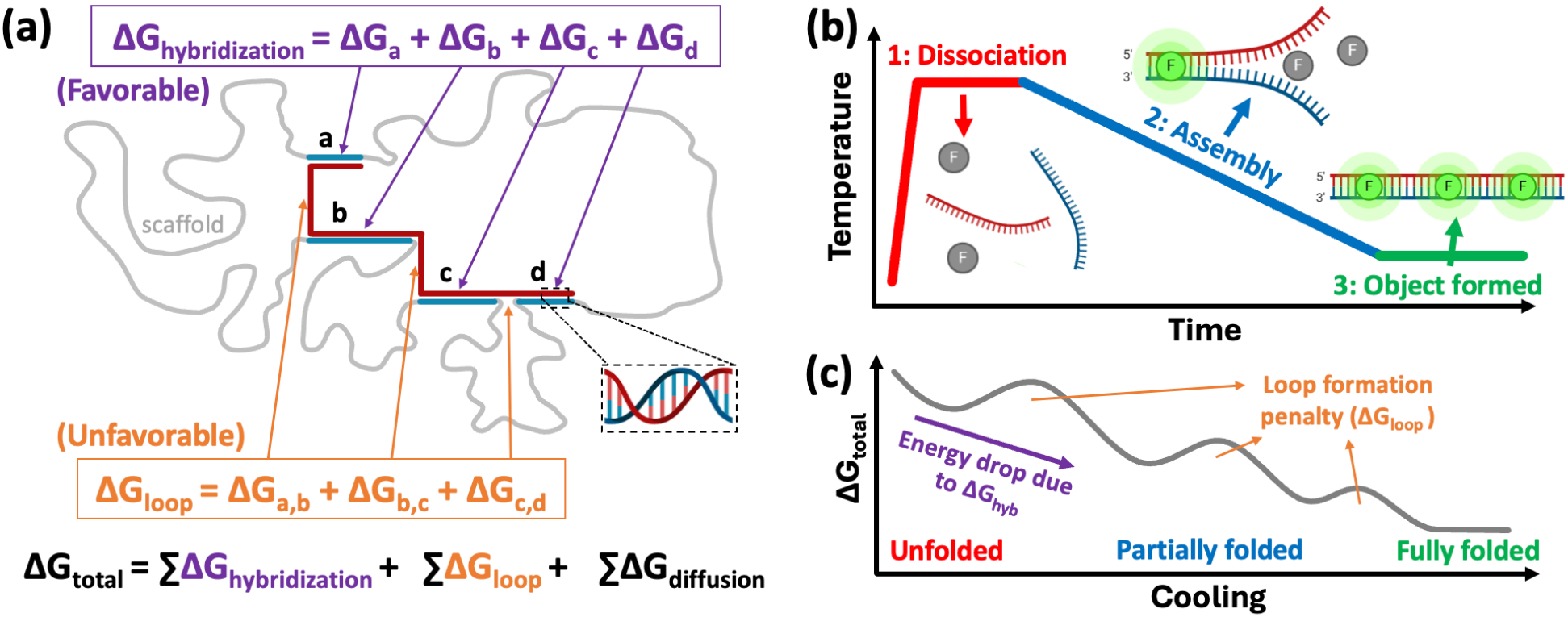
Free energy changes in DNA origami folding. a) Total free energy changes during the assembly process as the result of the interactions between staples and sections of scaffold by favorable hybridization (ΔG_hyb_), unfavorable entropic cost from loop formation (ΔG_loop_), and diffusion of staples. The schematic depicts the hybridization of one staple to different sections of the scaffold labeled a, b, c and d, inspired by the model from Aksel *et al*.^9^ b) Characterizing the assembly of DNA origami by real-time fluorometry involves 1) heating to dissociate all base pairs, 2) slowly cooling to allow for equilibrium assembly, resulting in 3) the final object properly formed. The fluorescence agent (F) attaches and fluoresces only to double-stranded DNA, signifying formation. (c) Total free energy changes in the system during assembly, where the fully assembled structure has a lower total free energy than the unpaired scaffold and staple DNA as the result of hybridization. The system should overcome loop formation penalty barriers during assembly, wherein the degree of such increase in energy would be lower in later peaks due to the cooperativity effect.

However, these thermodynamic terms are not the only parameters that govern the folding behavior of DNA origami structures, leading to discrepancies between computational models and experimental studies of folding. Namely, DNA origami folding is a highly cooperative process, often with a nucleation barrier. During folding, the lowest energy staple binding domains should bind first, mainly determined by length and sequence (*e.g.* ΔG_b_ in Figure 1a). Once a second binding domain from the same staple is bound (*e.g.* ΔG_a_ and ΔG_b_), closing a scaffold loop, the adjacent staple bridging the same two scaffold sections will have a facilitated binding event due to the reduced entropy penalty (*e.g.* Δ G_a,b_) necessary for binding. This event will facilitate cooperative assembly of the remaining staples in this region, and in some cases, throughout the entire structure. For many DNA origami designs, a critical number of initial binding events must occur before stable assembly can continue, which is known as the nucleation barrier.^31^ When adjacent staples on the same helix coaxially stack or the number of staple connections between scaffold sections increases, they create strong cooperative behavior. This increases the activation energy for nucleation, meaning that initial binding events must happen together before assembly stabilizes.^25^ Contrary to folding, unfolding is mostly an independent occurrence and usually happens at higher temperatures than the temperature at which the structure fully assembles, resulting in a hysteresis dependent on design complexity.^32^

Still, the role of cooperativity has not been fully captured. The cumulative role of individual scaffold-staple interactions does not fully describe the resulting nanostructure folding behavior. Instead, folding is primarily driven by cooperative effects due to the successive reduction in entropic penalty as scaffold loops form, as we demonstrate in this work. This study analyzes the folding behavior of a diverse set of DNA origami structures to elucidate the impact geometric complexity and staple design on folding behavior. We explore the relationship between design and the extent of cooperativity in assembly and examine the impact on assembly yield and time. The structures include a 30 helix-bundle (hb), flexible and rigid 24 hbs, a 2 hb, a 3 arm-linkage and a flat single-layer Longhorn logo. In addition, scaffold design strategies for the same structure are compared. Experimental folding characterization is performed through gel electrophoresis, electron microscopy, and real-time fluorometry during thermal annealing to analyze the folding across various time points and temperatures. The computational platform recently introduced by Aksel *et al.*^9^ is implemented to calculate theoretical energy profiles and compared to experimental results.

## Results & Discussion

### Folding and unfolding behavior

Real-time fluorometry experiments revealed distinct folding patterns for all DNA origami structures. Each initiates, progresses, and completes assembly at unique temperatures, offering insights into the assembly process. DNA origami nanostructures are typically assembled by heating the components above the melting temperature of all staples, then gradually cooling the solution to properly anneal each staple to the respective scaffold sections to form the intended structure. In this work, we perform these annealing ramps via real-time fluorometry with SYBR Green dye added in solution. The dye will intercalate and start to fluoresce only once DNA hybridizes, providing a real-time readout of DNA origami folding (Figure 1b), as first demonstrated by Sobczack *et al.*.^8^ If heated again, the structure will disassemble, as indicated by a loss of fluorescence. Results from fluorometry experiments in Figure 2 a-f show the rate for folding (cooling) and unfolding (heating) at each ramp temperature for all nanostructures studied. This includes a 30 hb, rigid 24 hb, flexible 2-arm version of the 24 hb, 3-arm 8 hb (3 arm-linkage), single-layer 2D Longhorn logo, and a 1D 2 hb. Each peak indicates a steep transition in folding behavior, calculated from florescence intensity changes as the amount of hybridization in solution changes. Folding ranges and the start of rapid unfolding are compared in Figure 2g, showing a narrower folding range at higher temperatures for the simpler 2D and 1D structures. Agarose gel electrophoresis (Figure 2h) and transmission electron microscopy (TEM) (Figure 2 i-n) confirm nanostructure assembly. The thermal ramp rate was optimized for the proper assembly of all structures (Figure S1-3).

**Figure 2.**
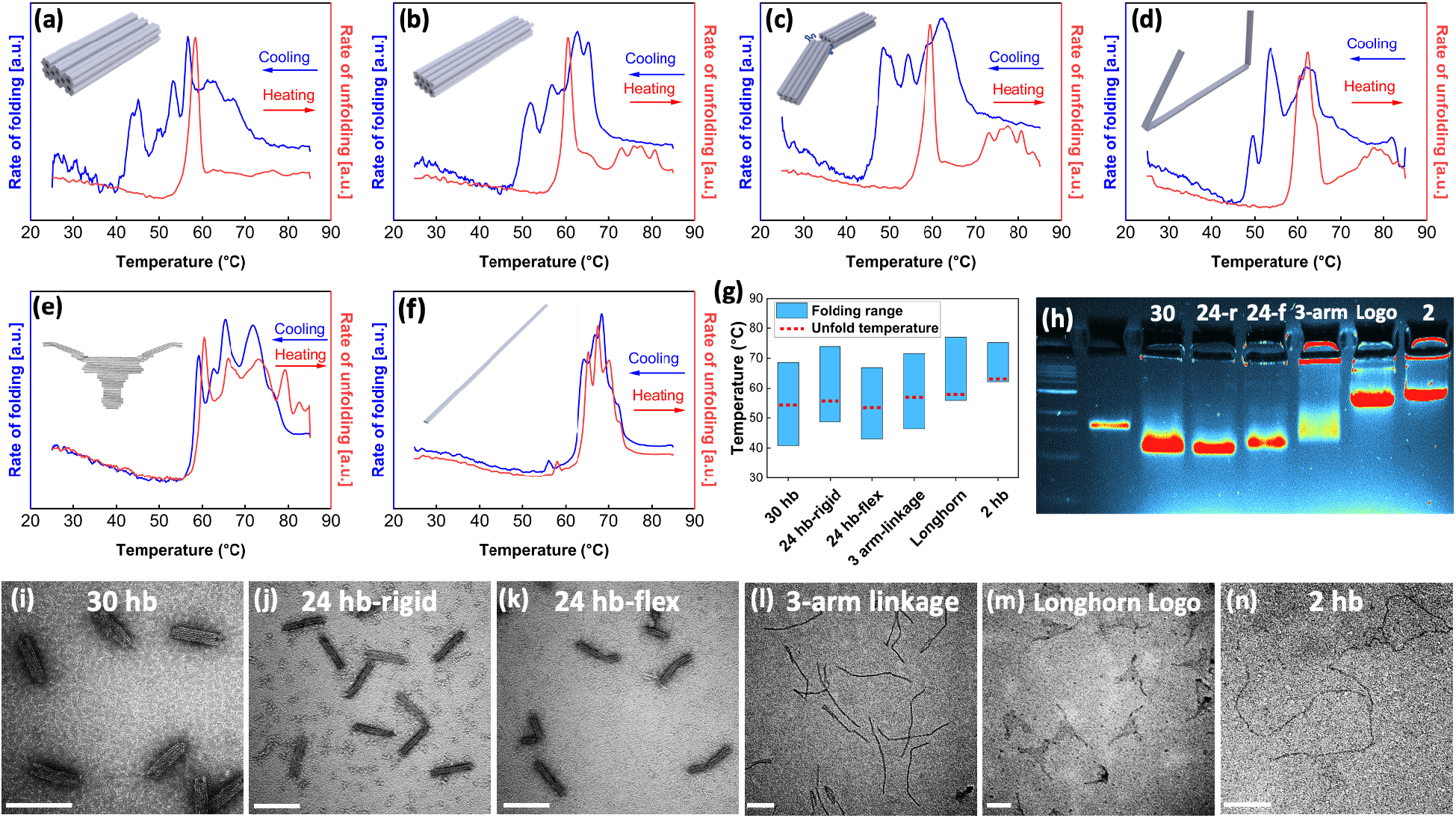
Folding and unfolding real-time fluorometry results of a) 30 hb, b) 24 hb-rigid, c) 24 hb-flex, d) 3-arm linkage, e) Longhorn logo, and f) 2 hb structures. Rates of folding and unfolding calculated from the change in fluorescence (hybridization) at each temperature. g) Folding range and start of unfolding temperatures from a)-f). h) Gel electrophoresis for each structure folded with the same thermal ramp as a)-f), indicating the successful formation of the intended structure. TEM images of folded structures: i) 30 hb, j) 24 hb-rigid, k) 24 hb-flex, l) 3-arm linkage, m) Longhorn logo, and n) 2 hb structures. Scale bars = 70 nm.

A key metric in folding analysis is the temperature at which folding is completed, T_fold_, which is seen here as the left side of the lowest temperature peak where no noticeable changes in folding are observed at temperatures below T_fold_.^8^ We observe drastic differences in T_fold_ as well as the number of distinct assembly steps between designs, where structure density and complexity result in lower T_fold_ and more folding steps. This is likely due to the greater entropic cost and embedded helices hindering folding of complex structures. For instance, in the case of the 30 hb (many densely packed helices), folding starts at around 68 °C and is completed at 41 °C with many folding steps. On the other hand, the 2 hb starts assembling at a higher 74 °C and completes the process at 61 °C, a narrow folding range of only 13 °C. Between these extremes, the 24 hb folds in a moderate 68-48 °C range. Adding complexity to the 24 hb with a single-stranded scaffold joint (24 hb-flex) lowers T_fold_ and adds more distinct folding steps due to the loss of cooperativity between arms, as discussed later. Full results are shown in Figure 2 and Table S1.

In a larger bundle of helices, there are more opportunities in the design to cross staples over to a neighboring helix, leading to a higher crossover density. The higher crossover density in the 30 hb created a nucleation barrier that requires cooperative binding of staples to initiate folding, whereas the 2 hb lacks a significant nucleation barrier, leading to its rapid and stochastic folding at higher temperatures.^25^ The presence of a nucleation barrier lowers T_fold_, as seen in the 30 hb. As temperature (*T*) is lowered during the annealing process, the decreased thermal fluctuations and contribution of entropy helps overcome the nucleation barrier for the 30 hb, delaying formation and lowering T_fold_ (Eq. 1).

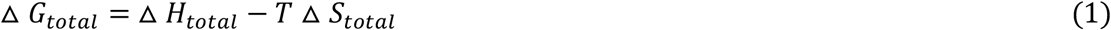

Contrary to the folding process, during unfolding, staple dissociation is not dependent on other staples. This phenomenon is mostly governed by the average penalty in the free energy of the object, leading to a sharp peak in the rate of unfolding, usually with a higher unfolding temperature (T_unfold_) relative to T_fold_. Staple binding order does not play a significant role in unfolding,^32,33^ as it does in folding. Once the structure is formed, the average hybridization energy is the same for all structures with similar scaffolds and staple design strategies. The average free energy penalty determines the unfolding process, meaning structures with higher penalty energy, such as the 30 hb, will unfold at lower temperatures. T_fold_, T_unfold_, and the difference between them are summarized in Figure 2g and Table S1 for each structure. We find that the additional complexity of structures with many helices or disjointed sections result in more hysteresis. Minimal hysteresis was seen in simple designs like the 1D 2 hb and 2D Longhorn structures. The presence of a nucleation barrier in the 3D designs is the primary reason for hysteresis, where unfolding and folding follow different pathways. Complex structures with a higher nucleation barrier (*e.g.* 30 hb) exhibit a more significant hysteresis than simple designs (*e.g.* 2 hb) with minimal nucleation barrier.

Differences in structure complexity (*e.g.* 30 hb vs 2 hb) influence many scaffold-staple design factors. The higher crossover density for structures like the 30 hb means each scaffold-staple pairing domain between crossovers will be shorter, resulting in a higher and less favorable ΔG_hyb_. More helices and crossovers also means looping more scaffold sections, raising ΔG_loop_. For instance, the 2 hb has potential staple crossovers every 21 bp, compared to every 7 bp in the 30 hb design, resulting in a consistently ordered staple design where almost every staple is nearly identical in length and crossovers, differing only by sequence (Figure S7f). Such a consistent ordered staple design leads to narrow total free energy change (Figure 3). The 30 hb has far more heterogenous staple lengths and crossover patterns and thus has greater free energy changes during assembly. The 2 hb also has only 2 scaffold crossovers, compared to 58 on the 30 hb, contributing to the differences in ΔG_loop_. This result is directly seen in folding and unfolding curves in Figure 2 where the 2 hb has a narrow folding range, 21.4 °C higher T_fold_, and minimal hysteresis, compared to the 30 hb structure.

**Figure 3.**
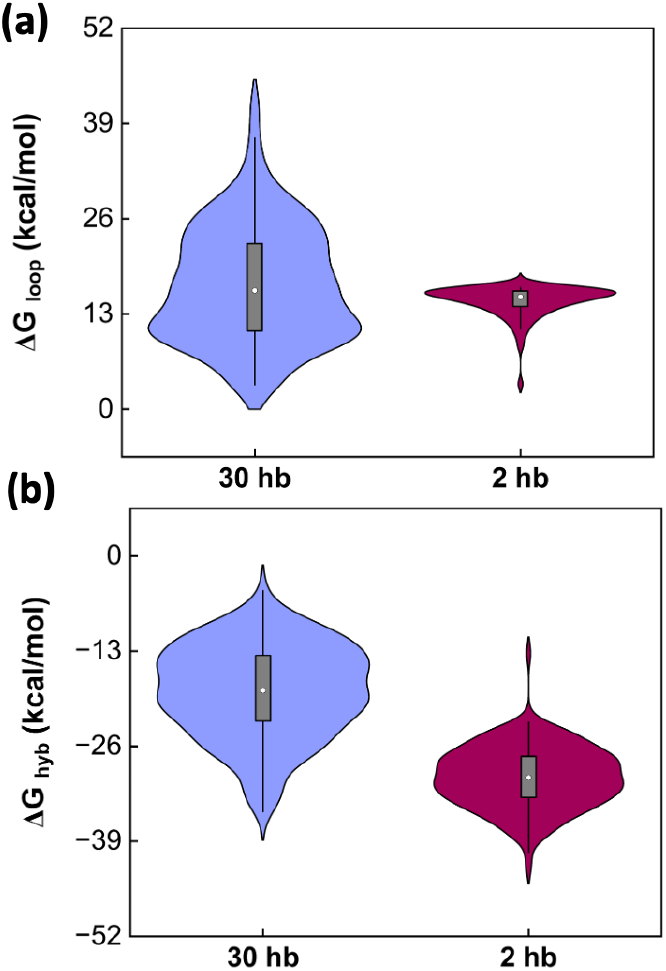
Theoretical energy contributions for every staple in the 30 hb and 2hb structures. a) ΔG_loop_ and b) ΔG_hyb_ show narrower energy distributions for 2 hb structure compared to 30 hb due to a more ordered scaffold-staple design, leading to smaller width of folding range.

### Impact of staple routing on folding behavior

The dynamic nature of the free energy of each staple plays an important role in folding behavior. Staple energies change throughout the folding process as the system changes and can be directed through scaffold-staple design.^23^ According to thermodynamic terms discussed in the introduction, increasing the length of staples and decreasing the number of crossovers can lead to a lower total free energy change, hence facilitating binding events as demonstrated with the results in Figure 3. However, achieving both factors and maintaining the structural integrity of the object is difficult, especially in complex designs. In this regard, to evaluate the influence of staple length and the number of crossovers, staple length was increased for the 3 arm-linkage at the cost of more crossovers per staple, as depicted in Figure 4a-b. In these designs, the circled staple breaks/nicks were removed throughout the structure, resulting in longer staple binding sections and longer overall staples. In total, 37 nicks were removed in the modified design with no other changes. Agarose gel electrophoresis (Figure 4c) and TEM images (Figure S4) showed that the design with longer staples demonstrated a significant drop in yield, from 94% to 20%, despite the addition of longer binding domains. The free energy changes of loop formation and hybridization of each staple are plotted in Figure 4 d-e. Although the longer staple design noticeably lowers ΔG_hyb_, the same staples resulted in higher ΔG_loop_ due to the 5 crossovers necessary for proper binding of the highlighted staples, compared to 2-3 crossovers per staple in the original design. This means the long staple design requires the closing of five scaffold loops per staple. The result is a higher entropy penalty shown here and likely kinetic traps during folding that hinder consecutive staple pairing. Therefore, we see further evidence that the entropy cost plays the larger role in the assembly behavior compared to the energy of hybridization.

**Figure 4.**
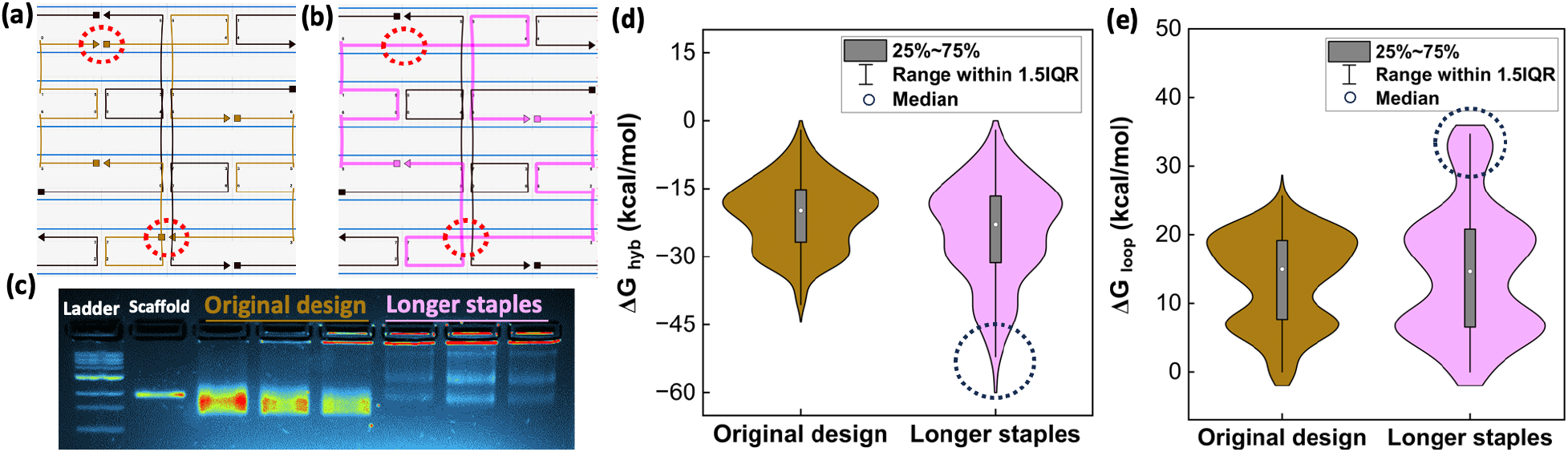
Effect of staple design on folding. Scaffold-staple designs for the 3 arm-linkage from cadnano show (a) before and (b) after increasing the length of staples by bridging the circled nicks. Pink staples are the longer modified staples, brown are the original staples, black staples are unchanged, and blue is scaffold. (c) Gel electrophoresis of both designs shows proper folding with the original design, and a significant decrease in yield for the modified design. Free energy changes for all staples show the modified staples have more favorable ΔG_hyb_ (d) due to the longer binding domains, but less favorable ΔG_loop_ (e) due to the increased number of crossovers per stable. The calculated total free energy change between designs is near zero, but experiments show the entropic penalty had a larger impact and entirely disrupts folding.

### The role of cooperativity

Cooperativity is present in both intrastrand (in the subsequence of the same strand) and interstrand (between different strands) folding.^34^ By increasing the length of a staple, intrastrand cooperativity is improved where initial base pairs help remaining base pairs by reducing their entropic penalty. However, our results in Figure 4 show that this also leads to a significant increase in entropy cost by increasing the number of crossovers per staple. The increased intrastrand cooperativity is not significant enough to enhance folding yield, where the higher entropy cost from the longer staple design deteriorated the yield. This supports the findings from Schneider *et al.*^35^ that non-local cooperative effects govern assembly behavior.

Thus, it is the interstrand cooperativity that plays a critical role in folding process of DNA origami.^36^ As discussed, once a critical number of staples are bound, successive staple binding is facilitated and folding continues more rapidly than predicted. This parameter has been difficult to quantify in theoretical models due to its strong dependence on structure and scaffold-staple design, leading to missing information in even the best predictive folding models.^9^ Our results show that cooperativity plays a pronounced and essential role in the assembly of complex designs, in which overcoming the nucleation barrier without the cooperative folding of staples for nucleation would be difficult. Figure 5 shows histograms of theoretical folding temperatures of all staples in each structure, as calculated using the recent model by Aksel *et al.*.^9^ This range is compared to the experimental T_fold_ of the structure, as measured through real-time fluorometry (Figure 2). Staples that folded at temperatures above their theoretical T_fold_ are shaded in blue. According to this analysis, at least half of the staples in most structures possess a lower theoretical T_fold_ than seen experimentally, which indicates the importance of cooperative folding. Our estimation of the effect of cooperativity is given in Table 1, calculated based on the percentage of staples that possess a theoretical T_fold_ lower than the experimental T_fold_.

**Table 1.** Comparing the total number of scaffold crossovers, number of internal scaffold crossovers not bridged by staples, and the contribution of cooperativity to the assembly of each structure.

| Structure | Total scaffold crossovers | Unbridged scaffold crossovers | Role of cooperativity (%) |
| --- | --- | --- | --- |
| Longhorn | 90 | - | 19.2 |
| 3-arm linkage | 30 | 16 | 50.0 |
| 24 hb-flex | 46 | 22 | 51.1 |
| 30 hb | 58 | - | 61.7 |
| 24 hb-rigid | 46 | - | 72.6 |
| 2 hb | 2 | - | 98.2 |

**Figure 5.**
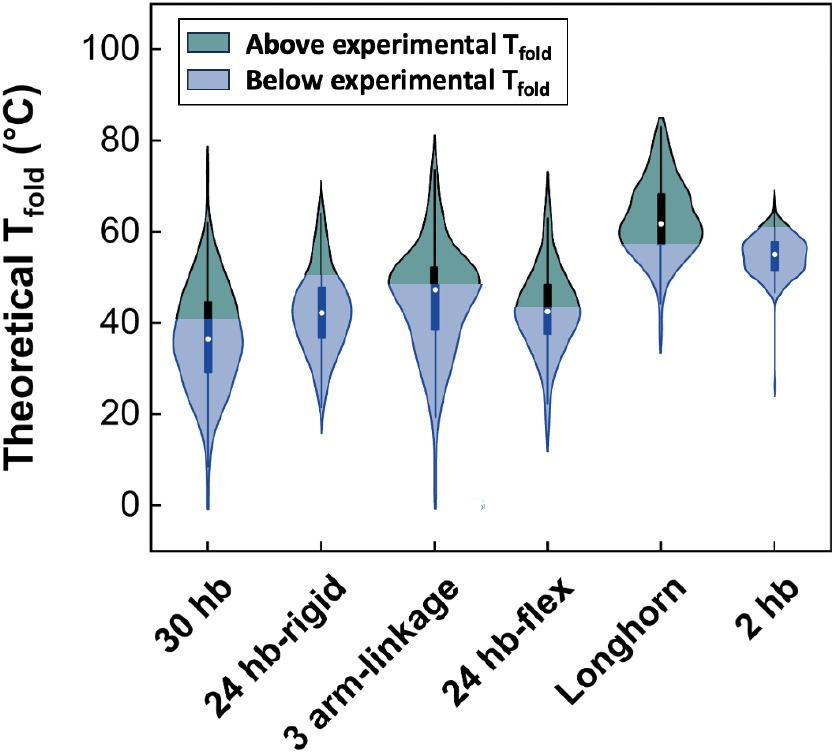
T_fold_ distributions for all staples in each structure calculated following Aksel *et al*.^9^ The blue/green boundary is the experimental T_fold_, meaning the staples in the blue region have a lower theoretical T_fold_ than experimental T_fold_. We attribute this difference to the cooperative effect during assembly. The dark rectangles represent the 25-75% percentile of the data, the bars show the 1.5 interquartile range, and the circle is the median.

As Table 1 shows, cooperativity in DNA origami structures is influenced by the number of scaffold crossovers, as each scaffold crossover presents an additional fold in the origami with potential to disrupt cooperative assembly. We describe cooperativity as the facilitation of neighboring staple binding. The more folds (crossovers) in the scaffold, the more likely it is to hinder this effect. This is seen as a consistent trend across all six structures. For example, the rigid 24 hb with 46 total scaffold crossovers is 72.6% cooperative, compared to the 61.7% cooperative 30 hb with 58 crossovers. At the extremes, we see the 2 hb with 98.2% cooperativity and 2 crossovers (1 fold of the scaffold) and the Longhorn with 19.2% cooperativity and 90 crossovers (∼45 folds). These 1D and 2D designs are expected to fold at higher temperatures compared to 3D designs. The theoretical T_fold_ (lowest folding temperature of any staple) is highest in the Longhorn, but due to the significant cooperative effects, we see the highest experimental T_fold_ in the 2 hb, as shown by the green-blue boundary in Figure 5 and Table S1. Beyond the total number of scaffold crossovers, their connection via bridging staples also plays a crucial role. For rigid bundles, internal crossovers are bridged with staples, but removing these bridging staples in multi-arm designs completely disrupts cooperative folding, limiting the effect to one arm instead of the entire structure. In structures like 24 hb-flex and the 3-arm linkage, the absence of bridging staples noticeably decreases the extent cooperativity. The 1-arm (rigid) and 2-arm (flex) versions of the 24 hb have identical scaffold designs with 46 crossovers, but the flexible version is 21.5% less cooperative. Structure design details are available in Figure S7.

To further examine the role of cooperativity in DNA origami assembly, a 24 hb structure was designed with two variations: one rigid bundle and one flexible (flex) version with a scaffold-connected joint between 2 rigid arms (Figure 6a). Both structures share an identical scaffold-staple design, except the flex version excludes 22 central staples across internal scaffold crossovers, while the rigid version retained them (Figure S7 b-c), inspired by previous work^37^. Figure 6b shows the theoretical T_fold_ for all staples. ΔG_loop_ and ΔG_hyb_ plots (Figure 6c-d) indicate that the staple free energies for both versions are nearly identical with the exception of the middle bridging staples. Although the bridging staples depict low ΔG_hyb_ (favorable), they introduce higher ΔG_loop_ (unfavorable) by adding crossovers bridging the two arms and higher ΔG_diffusion_ (unfavorable). The result of these subtle changes in theoretical folding behavior is negligible, with a total structure T_fold_ of 42.7 °C for the rigid 24 hb and 42.8 °C for the flexible version, as depicted in Figure 6e. However, despite this theoretical similarity, our experimental results reveal a T_fold_ of 48.7 and 43.0 °C for the rigid and flexible 24 hb structures, respectively (Figure 6e-f). This tells us that the degree of cooperativity is lower in the flex version than in the rigid version, as shown with full thermal ramps. Connecting the two halves in the rigid version extends the cooperative effect throughout the entire structure, facilitating formation of the other arm, compared to the 2-arm flexible version which limits the cooperative enhancement to a single arm.

**Figure 6.**
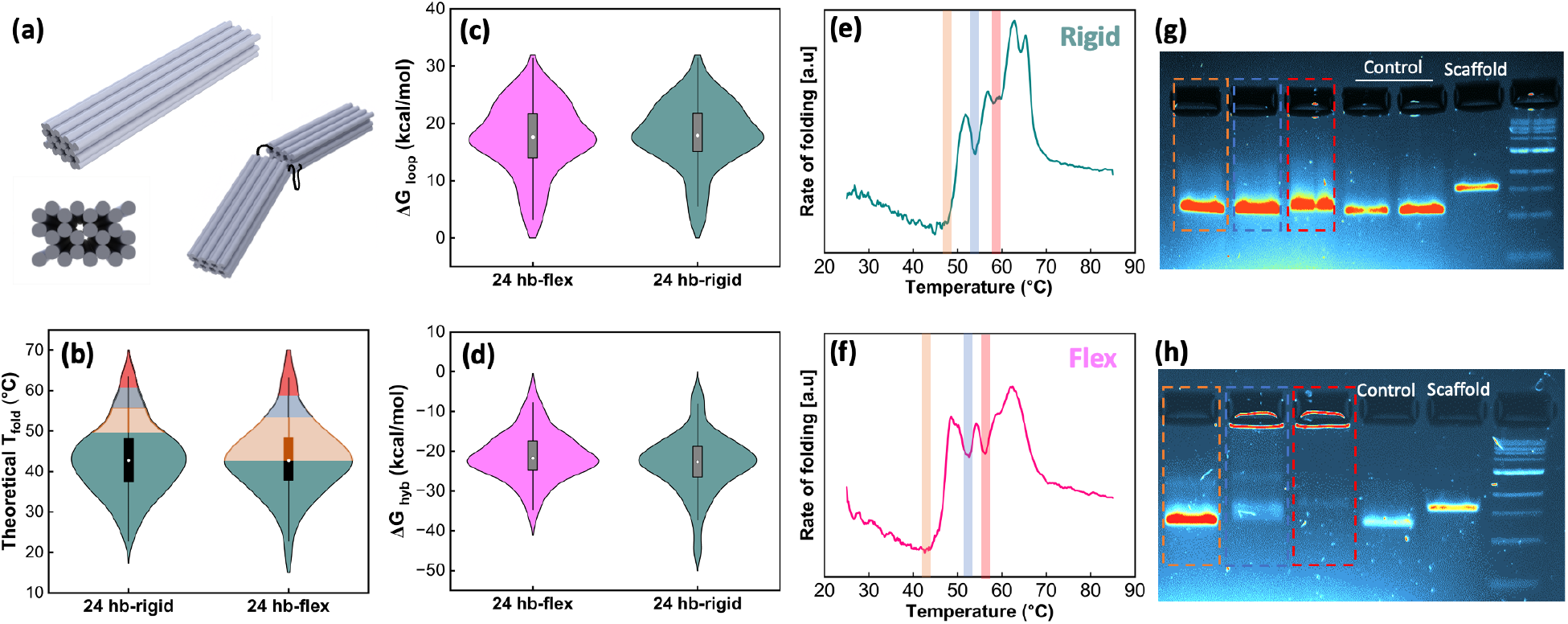
The impact of unbridging on cooperative assembly. (a) 24 hb rigid and flex structures have identical designs except 22 staples that bridge internal scaffold crossovers are removed in the flex version. Theoretical distributions of (b) T_fold_, (c) ΔG_loop_ and (d) ΔG_hyb_ for all staples, calculated using code from Aksel *et al*.^9^ show nearly identical energies between rigid and flex versions. Real-time fluorometry curves of e) 24 hb-rigid and f) 24 hb-flex objects with three local energy minima highlighted in red, blue, and orange. Agarose gel electrophoresis analysis of g) 24 hb-rigid and h) 24 hb-flex frozen at the highlighted temperatures during the annealing ramp. Dotted boxes in gels (g-h) and shaded regions in (b) correspond to corresponding temperatures on fluorometry curves (e-f). Results show more cooperative folding in the rigid version. In (b-d), the dark rectangles represent the 25-75% percentile of the data, the bars show the 1.5 interquartile range, and the circle is the median.

To investigate these differences more thoroughly, structures were shock-frozen at the temperatures highlighted in Figure 6e-f, where a dip in the rate of folding is seen. Gel electrophoresis analysis in Figure 6g-h shows the extent of folding at each temperature, showing near complete folding for the rigid, but almost nothing resembling folded structures at intermediate temperatures for the flex design. Control bands show properly folded structures. For comparison, theoretical T_fold_ is shown in Figure 6b with each temperature range highlighted, again showing a large discrepancy between theoretical and experimental T_fold_ due to the role of cooperativity. This is apparent with the intermediate folding analysis and higher T_fold_ of the rigid structure, indicating a more robust and faster assembly with the majority of folding occurring during the first ∼60 °C folding step, well above the theoretical T_fold_.

It has been reported that DNA origami structures can be folded at constant temperatures of their T_fold_.^38–40^ To measure their robustness, both versions of the 24 hb were heated at constant temperatures of 50 and 55 °C for 30 minutes, intermediate ranges above their T_fold_, and their gel electrophoresis analysis is shown in Figure 7a. Whereas the rigid structure formed at both temperatures, the flex design only assembled at 50 °C, suggesting a less robust and less cooperative design. Based on the annealing profiles (Figure 7b), at 55 °C the rigid structure was farther along in the folding process than the flex structure and folding was likely able to complete with enough time due to cooperative effects, even at a temperature higher than T_fold_. Furthermore, to study their constant temperature folding kinetics, the 30 hb, 24 hb-rigid, and 24 hb-flex structures were heated at their T_fold_ (Table S1) for 0, 0.5, 1, 5, 10 and 30 minutes (gel electrophoresis images shown in Figure 7 c-e). The required time to properly assemble each structure with near 100% yield was 30, 5, and 1 minute for the 30 hb, 24 hb-flex, and 24 hb-rigid, respectively. Easier folding is apparent for a simpler cross section with fewer embedded helices (24 hb vs. 30 hb) and for a continuous structure compared to a jointed 2-arm bundle. Thus, folding kinetics appear to be influenced more by structure design than cooperativity. Folding yield and migration distance metrics for these gels are provided in Table S2.

**Figure 7.**
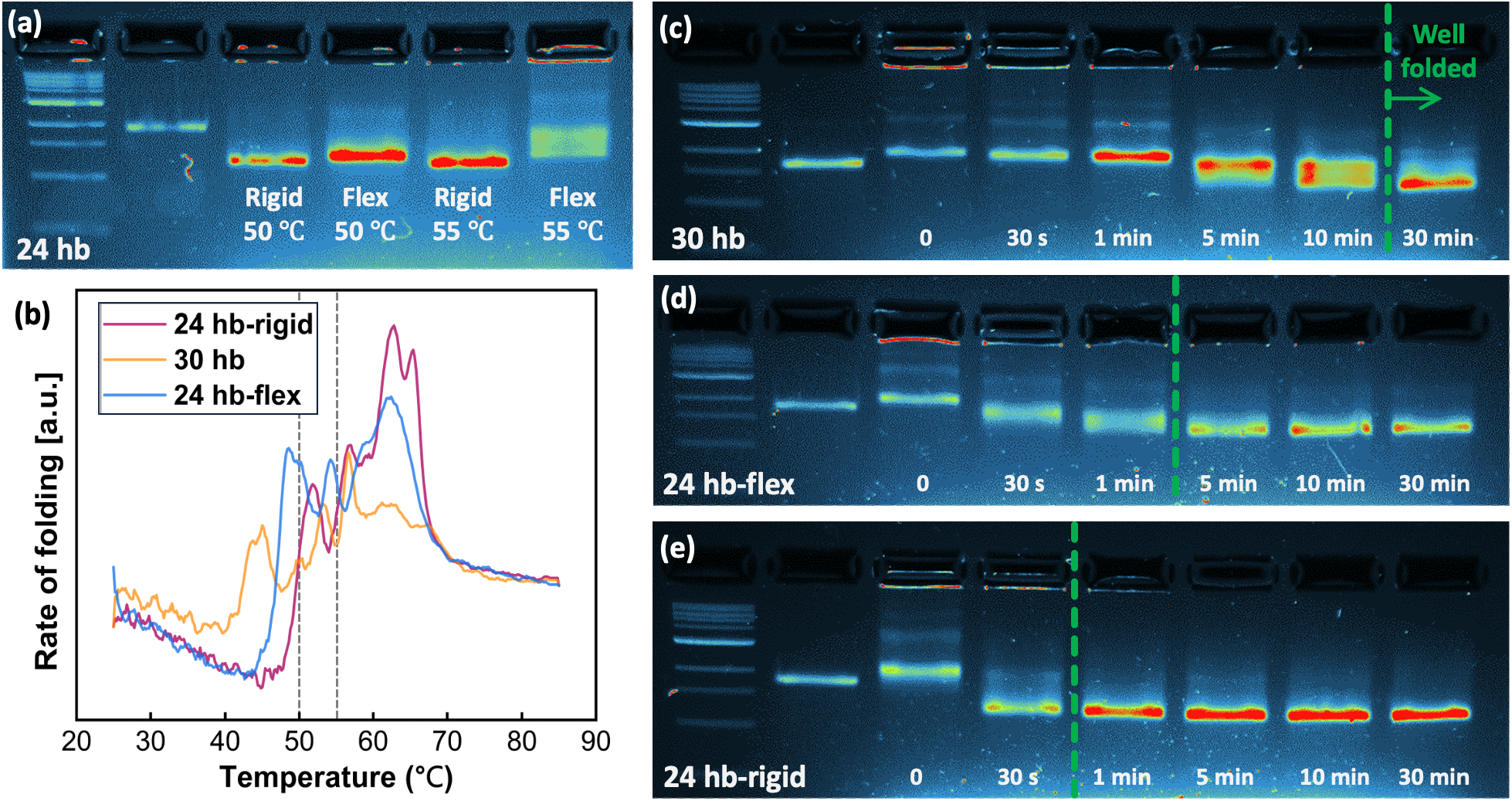
Kinetics of constant-temperature folding at T_fold_. a) Gel electrophoresis analysis of 24 hb rigid and flex objects folding at constant 50 °C and 55 °C for 30 min. (b) Real-time fluorometry results for the three structures studied here. Gel electrophoresis was used to evaluating folding of c) 30 hb, d) 24 hb-flex and e) 24 hb-rigid objects after heating at their T_fold_, (41, 43 and 47 °C, respectively) for the times indicated. Proper folding is indicated by the green dashed line with well-folded structures to the right of the line. The first and second lanes of all gel images are ladder and 7249-scaffold, respectively.

DNA origami structures with more complicated designs (*e.g.*, more crossovers per staple and higher ΔG_loop_) as well as structures with higher nucleation barriers that rely more on cooperativity, require longer times for nucleation and complete assembly when annealed at constant temperatures.^25,33^ Consequently, in the case of a complex design, the probability of rapid and successful pairing of each staple may be lower or slower, as the staples will require lower temperatures and/or longer times to anneal due to the larger nucleation barrier, hence, diminishing or delaying the “helper” ability for subsequent staple pairing. This is apparent with the higher nucleation barrier in the 24 hb flex and 30 hb structures that demands a stronger cooperative effect and results in a longer annealing time necessary for efficient folding. The cooperativity influence is terminated when staples do not maintain coaxial stacking to reduce entropy cost for neighboring staples, such as the disjointed 24 hb flex design. This leads to lower T_fold_ and slower folding.

### Strategies to enhance folding yield

From our results presented here and the cumulative literature we have discussed, we identify many factors that affect folding yield of DNA origami. Scaffold-staple design and thermal ramp protocols have a more significant effect once the salt concentration is optimized. Regarding the scaffold-staple design, the energy changes of each staple during binding events should be considered, and the trade-off between the energy of hybridization and loop formation should be optimized, where the latter has shown a more pronounced impact. Concerning the thermal ramp, certain temperatures should be focused on or completely avoided to achieve the maximum yield. Real-time fluorometry is a valuable technique that can be utilized to fulfill this goal by determining the temperatures at which folding is initiated and completed. This is unique for every design and sensitive to even a few staple changes. These considerations are also crucial for multicomponent structures, where the sequence of folding sections matters^40,41^. Additionally, through a folding and unfolding process, this technique can assess whether the thermal ramp used resulted in the formation of the intended DNA origami structure or caused agglomeration and undesirable base pairing events, without the need for gel electrophoresis or TEM (Figure S1-2). Finally, in some cases, heating for long enough above the temperature where folding starts can reduce the folding yield or even prevent it due to increased undesirable aggregation (Figure S2 and S3).

By focusing on the folding temperature range, not only can the yield be improved, but the folding time can be noticeably lowered. Figure 8 compares the folding yield for thermal ramps of 2.5 days, a 1 h constant ramp of the full temperature range (same as Figure 2), and a ramp focused on the folding temperature range (1.6-2.5 h). The associated gel electrophoresis images are shown in Figure S6. For the focused ramp, structures were annealed only in their folding range, decreasing the temperature by 0.1 °C every 30 seconds. The 2.5-day and 1-hour ramps covered the full temperature range from 65-25 °C. The results in Figure 8 show that the focused ramp achieved equal or better folding yield than the 2.5 day or rapid full range ramps for all structures by avoiding aggregation and kinetic traps at unnecessary temperature ranges. For instance, the folding yield of the Longhorn and 3-arm linkage were enhanced by about 17%, compared to the 2.5-day ramp, and the latter increased yield 22.7% compared to the rapid 1-h ramp. Performing one real-time fluorometry folding experiment to determine the optimal folding range, then using this strategy for enhanced rapid folding can save significant time while maximizing folding yield.

**Figure 8.**
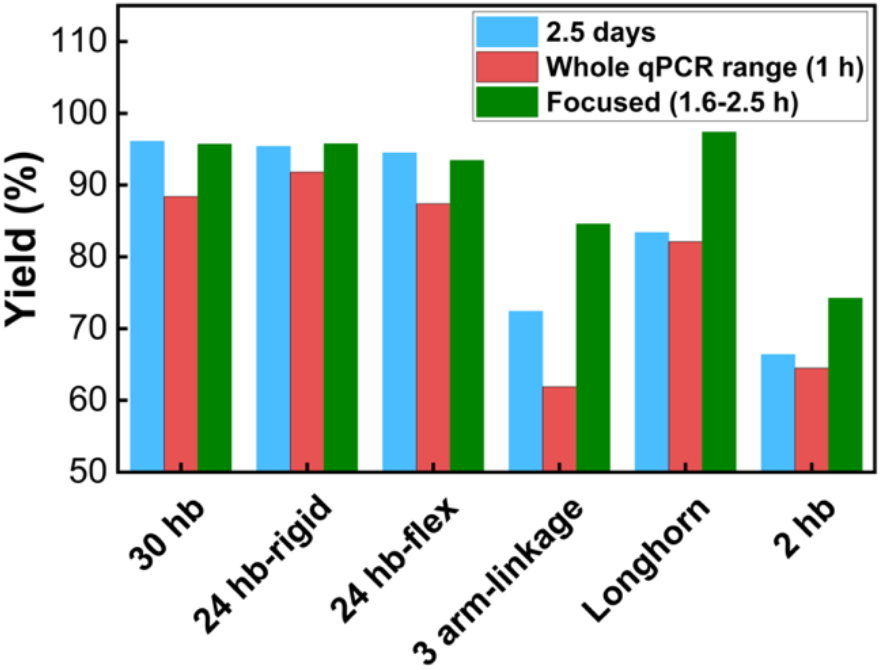
Folding yield analysis of all structures after thermal annealing ramps of 2.5 days, the whole real-time fluorometry range used in Figure 2 (1 h total), and new focused thermal ramps covering only the temperatures in the folding range of the structure (Figure 2g) at 5 min/°C. The corresponding gel electrophoresis images are shown in Figure S6.

We have established two important lessons to enhance DNA origami folding yield: 1) in the design phase, ΔG_loop_ is of paramount importance and should be minimized while preserving the design integrity, and 2) real-time fluorometry is a valuable tool that gives insight into the folding temperature range of the intended design, and thus, an optimized thermal ramp can be designed to focus on this range. In addition, to develop a multicomponent device, cooperativity and nucleation barriers can be tailored for directed assembly.

## Conclusions

This study examined the folding behavior of DNA origami, revealing the impact of design parameters on free energy changes, cooperativity, folding yield, and folding kinetics. Real-time fluorometry analysis showed that structures with more intricate designs (*e.g.,* embedded helices or jointed arms) exhibited lower T_fold_ and multiple folding steps. Energy analysis of 3-arm linkages indicated that loop formation energy (entropy) plays a larger role than hybridization energy, with long staples hindering folding due to increased entropy cost, despite more favorable binding domains. We show that cooperativity in DNA origami folding is strongly influenced by the number of scaffold crossovers and their potential connections via bridging staples. In rigid structures, fewer scaffold crossovers enhanced cooperativity, ranging from 98% cooperative with 2 crossovers to 19% cooperative with 90 crossovers. Removing bridging staples between internal scaffold crossovers, which creates multi-arm structures, was shown to decrease cooperativity by over 20% in the 24 hb, while lowering T_fold_ and slowing assembly. Quantifying the control of cooperativity and its impact on folding have been missing in previous experimental and computational studies. Building from the recent computational models from Aksel *et al.*,^9^ our experimental results can be used to improve yield and assembly time of complex DNA origami nanostructures. This is demonstrated with focused annealing ramps based on new insights from fluorometry results that enhance yield up to 17% while cutting the required assembly time to just 1-2 hours. This impact was especially noticeable for complex structures like the 3-arm linkage by focusing on the necessary temperatures and preventing aggregation at unnecessary temperatures. Optimizing free energy contributions and the extent of cooperativity in the design phase are essential for overcoming cost, yield, and efficiency limitations in large and complex DNA origami assemblies.

## Materials and methods

### DNA origami designs and fabrication

DNA origami computer-aided design software caDNAno^42^ was employed to design all structures (Figure S7). Finite element modeling tool CanDo^43,44^ was also used to analyze and inform designs. In addition, a recently introduced computational tool^9^ was used to calculate the energy profiles. The structures were fabricated using 7249-nucleotide (30hb, 24hb-rigid, 24hb-flex, 2hb) and 8064-nucleotide (3 arm linkage, Longhorn logo) scaffolds (Tibilit Co., Germany and Bayou Biolabs) and engineered staple oligos ordered were from Integrated DNA Technology. For the self-assembly process of all studied structures, 20 nM scaffold with the associated 500 nM staples were mixed in folding tubes, including 50 mM Tris, 10 mM EDTA and 18 mM MgCl_2_, similar to the process employed by Castro *et al*..^44^ This was followed by a thermal annealing ramp where the temperature decreased from 85 °C to 25 °C by -0.2 °C and heating at each temperature for 13 seconds. In addition, for the focused thermal ramps, structures were heated in their folding temperature range where data was obtained from real-time fluorometry results. Here, the temperature was decreased by -0.1 °C and held at each temperature for 30 seconds. Also, for constant temperature annealing, structures were held at the corresponding temperatures for the studied time points. For comparison, thermal annealing was conducted over 2.5 days, starting at 65°C and cooling to room temperature with a gradual decrease of 1 °C per step. The sample was held at each temperature for 1 hour until reaching 62 °C, 2 hours until 59 °C, 3 hours until 46 °C, 1 hour until 40 °C, and finally 30 minutes until 25 °C.

### Structure characterization using gel electrophoresis and TEM

To confirm the folding efficiency, the reaction products were mixed with a homemade loading dye electrophoresed on gels containing 2% agarose in 0.5x TBE with 11 mM MgCl_2_ and 1 μM SYBR safe for 1.30 h at 70 V in an ice bath. The Analytik Jena imager (Jena, Germany) was used to visualize the gels, and images were processed with VisionWorks software to calculate yield. The fast-leading band for structures in different wells were cut and purified utilizing freeze and squeeze DNA gel extraction spin columns (Bio-rad, Hercules, CA). For TEM imaging, 5 µL of purified objects were deposited and incubated for 1 min on glow-discharged formvar-supported carbon-coated TEM grids (Electron Microscopy Sciences, Hatfield, PA) and stained utilizing 1% uranyl acetate. Images were taken at 80 kV acceleration voltage with a FEI Tecnai G2 Spirit TEM.

### Real-time fluorometry

DNA origami objects were studied using a real-time PCR system (Bio-rad, CFX Opus 96) with the thermal ramp discussed in the fabrication section. In this experiment, the fluorescence intensity changes for the DNA origami structures were subtracted from their corresponding staple-only samples, and the derivative of the subtracted value that was denoted as the ‘rate of folding’ was plotted against temperature. Similar to the folding studies, unfolding analysis was performed once the structures were fully formed by heating from 25 to 85 °C. The heating/cooling ramp was optimized by trying different ramps (Figure S1-3) and kept consistent for all studied structures to exclude its influence on the assembly process. This ramp was implemented by increasing/decreasing the temperature by 0.2 °C and heating/cooling at each temperature for 13 seconds.

## AUTHOR INFORMATION

## Corresponding Author

*Alexander E. Marras

## Author Contributions

The manuscript was written through contributions of all authors. All authors have given approval to the final version of the manuscript.

## Funding Sources

The conducted research was supported by The University of Texas at Austin and the National Institute of General Medical Sciences at the National Institutes of Health under award R35 GM154984.

The authors would like to thank Michelle Mikesh (Electron Microscopy Specialist, Sauer Structural Biology Laboratory at The University of Texas at Austin) for assistance with TEM.

## ABBREVIATIONS

### Supporting Information Available

The supporting information is available and contains: real time fluorometry results at different thermal ramps (folding and unfolding), gel electrophoresis images of studied different thermal ramps, TEM images of 3 arm-linkage with normal and longer staple sections, theoretical ΔG_loop_ and ΔG_hyb_ of all studied DNA origami structures, and cadnano designs.

