## Supplemental Information for "Unraveling the Folding Dynamics of DNA Origami Structures"

### Supplemental Methods

#### Controlled reactions

The temperature-dependent assembly of 24 hb-rigid and flex DNA origami structures was conducted based on their real-time fluorometry curves. For these experiments, 50 µL of reaction mixture was aliquoted into PCR tubes, heated to 85 °C, and then cooled following the same thermal ramp protocol used in Figure 2 (85 to 25 °C, -0.2 °C per step, with a 13-second hold at each temperature). Once the samples reached the target temperatures, they were removed from the thermal cycler and quenched in liquid nitrogen. For the constant-temperature annealing protocol, samples were heated to the desired temperature and maintained for a specified duration before being similarly quenched in liquid nitrogen. After preparation, all samples were thawed and promptly analyzed using gel electrophoresis or TEM.

#### Real-time fluorometry experiments

To monitor fluorescence intensity changes, SYBR Green was added to the folding tubes of various DNA origami structures at a ratio of one molecule per 900 DNA base pairs. Each DNA origami sample consisted of 12 µL of the same reaction components as the folding assembly, with the addition of SYBR Green, and was aliquoted into PCR tubes. Additionally, reference reactions were prepared, including (i) only the scaffold (excluding staples), (ii) only staples (excluding the scaffold), (iii) only buffer (excluding both scaffold and staples), and (iv) a DNA hairpin with a melting temperature of approximately 74 °C, which was analyzed using the following sequence:

5′-TCAACATCAGTCTGATAAGCTACACGAGATCAGACTGATGTTGA-3′

Different thermal ramping protocols were tested to optimize the cooling rate for maximizing the yield of all studied DNA origami structures while covering a broad temperature range to capture their assembly behavior. Temperature was decreased from 85 to 25 °C at rates of -0.1 °C per step with hold times of 17, 72, and 312 seconds, as well as -0.2 °C per step with a 13-second hold. Figure S1 presents the real-time fluorometry results for all tested ramps. To further assess the impact of ramping rates, samples were analyzed using 2% agarose gel electrophoresis after each thermal protocol (Figure S3). For such thermal ramp to cover the range of 85-25 ℃, the results indicate that slower cooling rates led to minimal structure formation across all designs, and increasing the cooling rate improved the yield. Notably, the -0.2 °C per step with a 13-second hold produced the highest yield for nearly all structures. For the focused thermal ramps, structures were heated only to the temperature at which fluorescence intensity began to increase, then gradually cooled at a rate of -0.1 °C per step, with a 13-second hold at each temperature, until the signal stabilized. For example, one such ramp spanned from 68 °C to 41 °C for 30 hb.

Unfolding investigations were conducted after the cooling ramp for all tested thermal protocols, using a heating rate identical to the cooling rate (Figure S2). The real-time fluorometry unfolding plots of samples that did not successfully form the desired DNA origami structures exhibited a unique feature: they were nearly identical and showed fluorescence intensity changes only at very high temperatures (~80 °C), suggesting aggregation dissociation. In contrast, the optimized thermal ramp, which led to well-formed DNA origami structures, displayed distinct unfolding behaviors for different designs. These findings indicate that real-time fluorometry can be used to assess whether a given thermal ramp successfully facilitates proper DNA origami folding or induces undesired interactions, such as scaffold-scaffold, staple-staple, or staple-scaffold associations.

**Table S1.** $T_{fold}$ and $T_{unfold}$ for each structures, their difference, and width of folding range temperature obtained from real-time fluorometry curves (Figure 2).

| Structure | $T_{fold}$ (℃) | $T_{unfold}$ (℃) | Hysteresis ($T_{fold}$-$T_{unfold}$) (℃) | Width of folding range (℃) |
|---|---|---|---|---|
| **2 hb** | 62.2 | 63.4 | 1.6 | 13.0 |
| **Longhorn** | 55.8 | 57.8 | 2.0 | 21.2 |
| **24 hb-rigid** | 48.7 | 55.8 | 6.3 | 18.4 |
| **24 hb-flex** | 43.0 | 53.0 | 10.0 | 23.9 |
| **3-arm linkage** | 46.4 | 56.6 | 10.2 | 25.2 |
| **30 hb** | 40.8 | 54.6 | 13.8 | 27.8 |

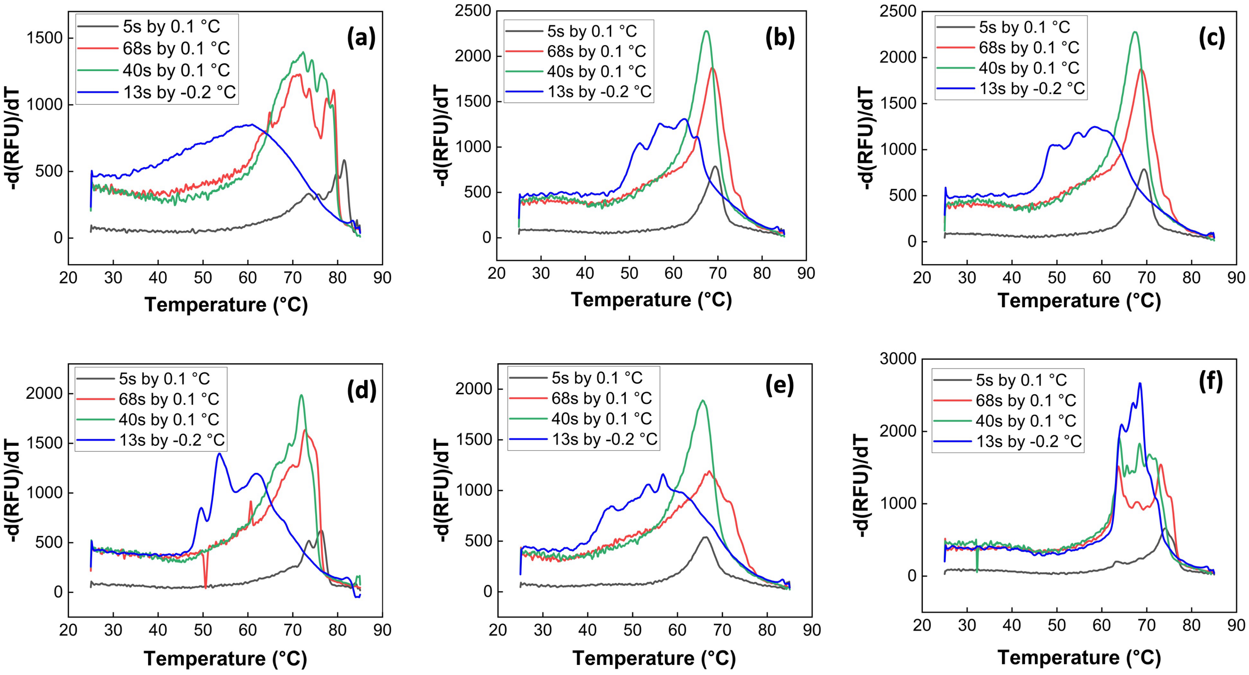


**Figure S1.** Real-time fluorometry results for (a) 30 hb, (b) 24 hb-rigid, (c) 24 hb-flex, (d) 3-arm linkage, (e) Longhorn, and (f) 2 hb, after annealing from 85 to 25 ℃ with different rates such as decreasing the temperature by 0.1 ℃ and staying at each temperature for 312 seconds (blue), 72 seconds (red), and 17 seconds (black) as well as dropping the temperature by –0.2 ℃ with 12 seconds time for each temperature during cooling (green). These results are before staple reduction from the DNA origami object data.

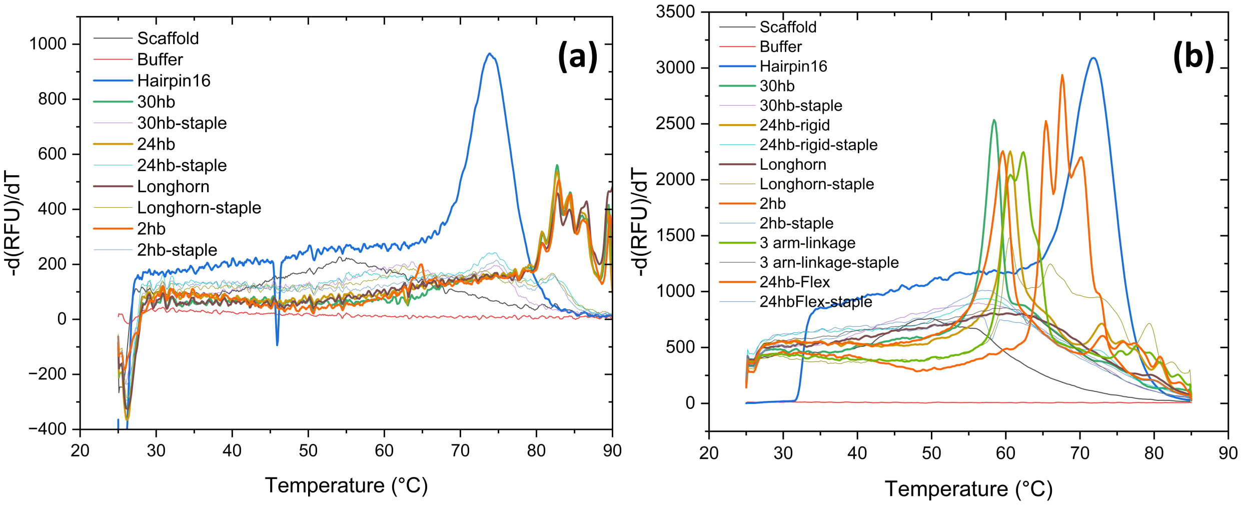


**Figure S2.** Unfolding real-time fluorometry result of different structures that were fold previously by cooling from 85 to 25 ℃ by (a) -0.1 ℃ for 312 seconds at each temperature (longest thermal ramp) and (b) -0.2 ℃ for 13 seconds, before staple reduction from DNA origami object.

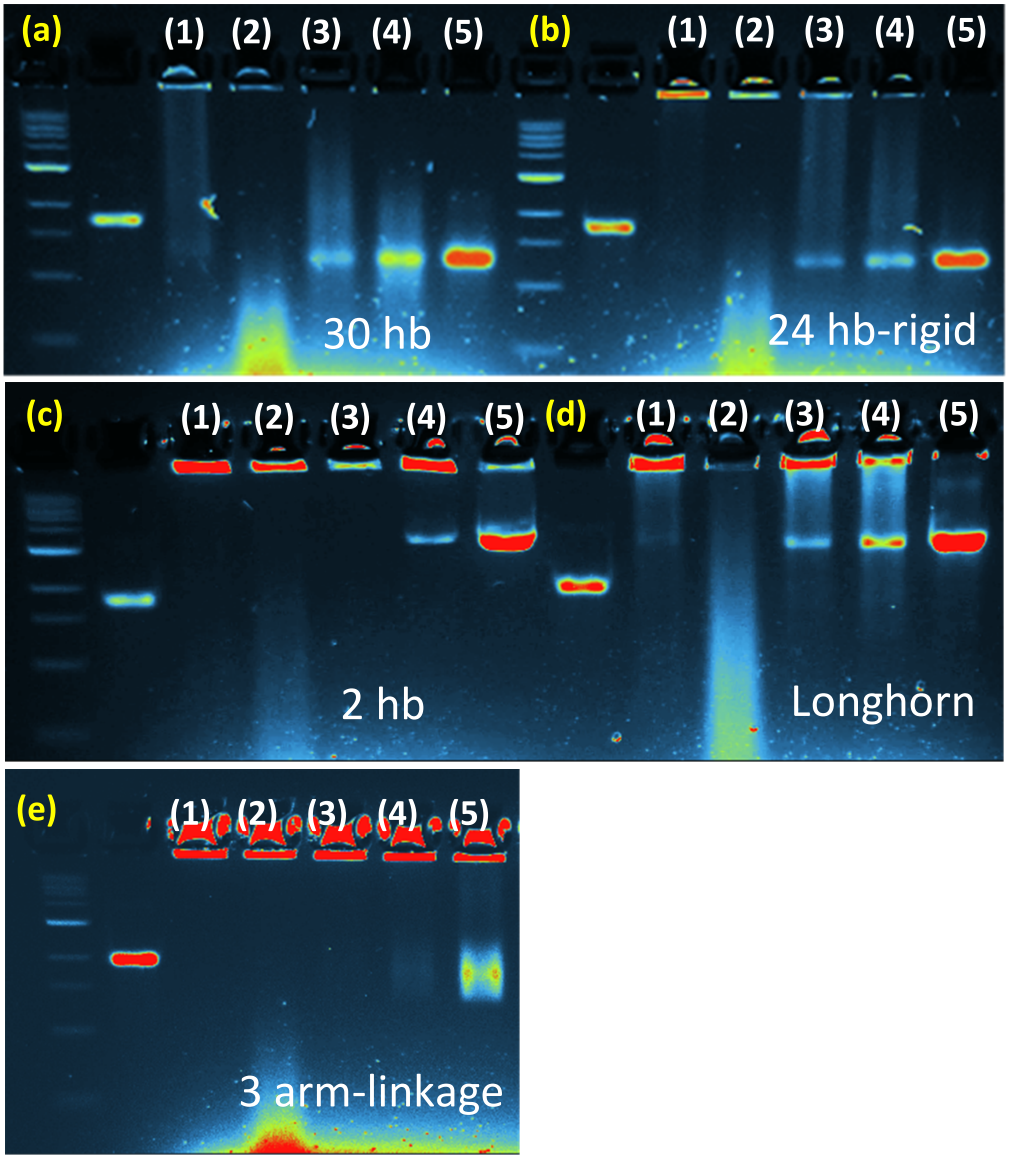


**Figure S3.** 2% Agarose gel electrophoresis of (a) 30 hb, (b) 24 hb-rigid, (c) 2 hb, (d) Longhorn and (e) 3 arm-linkage structures after cooling from 85 to 25 ℃ with decreasing the temperature by -0.1 ℃ and heating at each temperatures for (1) 312 seconds, (2) 72 seconds, (3) 17 seconds, and (4) temperature drop of -0.2 ℃ for 12 seconds. The first lane in all gel images is 1kbp ladder and the second lane is scaffold as the control.

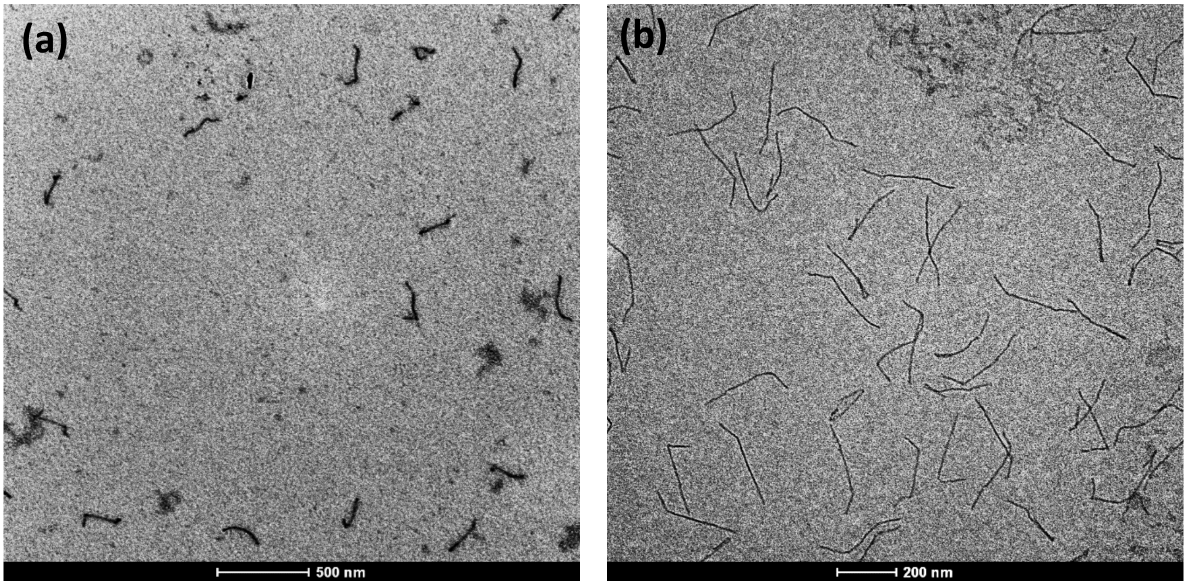


**Figure S4.** TEM images of 3 arm-linkage: (a) the design with longer sections of staples and (b) original design.

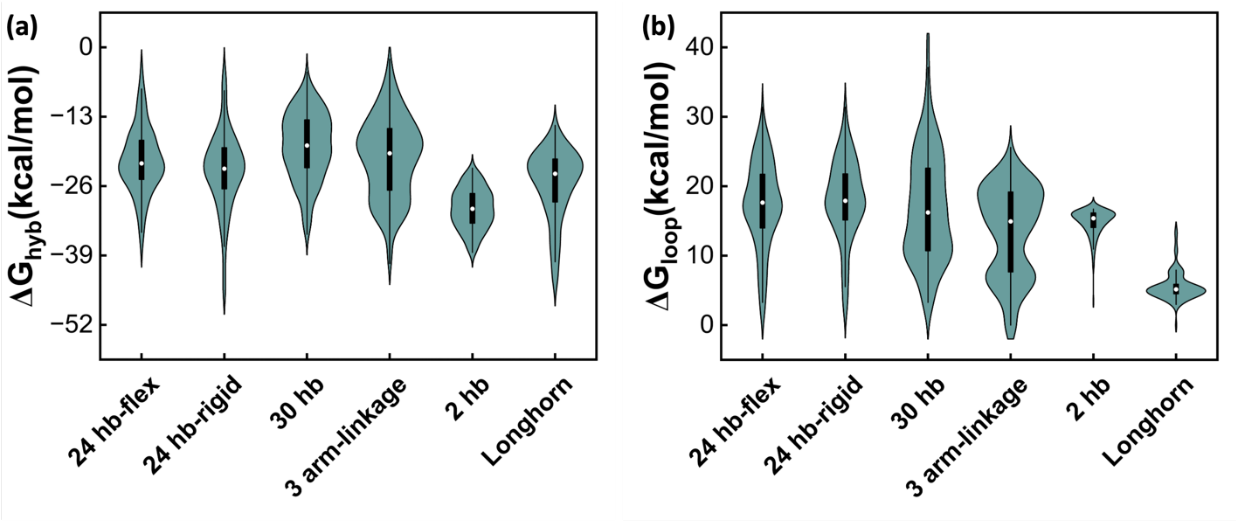


**Figure S5.** (a) Hybridization and (b) loop formation energies for all staples in all studied DNA origami structures obtained by using the computation method introduced in Ref[2].

**Table S2.** Folding yield and migration distance of the three studied designs in Figure 6

| Structure | Sample | Band migration (px) | Intensity (%) |
|---|---|---|---|
| **24 hb-flex** | 0 | 143.47 | 25.51 |
| | 30 s | 148.47 | 58.05 |
| | 1 min | 149.47 | 68.49 |
| | 5 min | 162.47 | 78.94 |
| | 10 min | 180.47 | 80.6 |
| | 30 min | 180.47 | 83.31 |
| **24 hb-rigid** | 0 | 130.85 | 63.52 |
| | 30 s | 174.85 | 69.88 |
| | 1 min | 181.85 | 84.24 |
| | 5 min | 179.85 | 87.59 |
| | 10 min | 182.85 | 96.5 |
| | 30 min | 187.85 | 97.86 |
| **30 hb** | 0 | 124.53 | 63.52 |
| | 30 s | 168.53 | 69.88 |
| | 1 min | 175.53 | 84.24 |
| | 5 min | 173.53 | 87.59 |
| | 10 min | 176.53 | 96.5 |
| | 30 min | 181.53 | 97.86 |

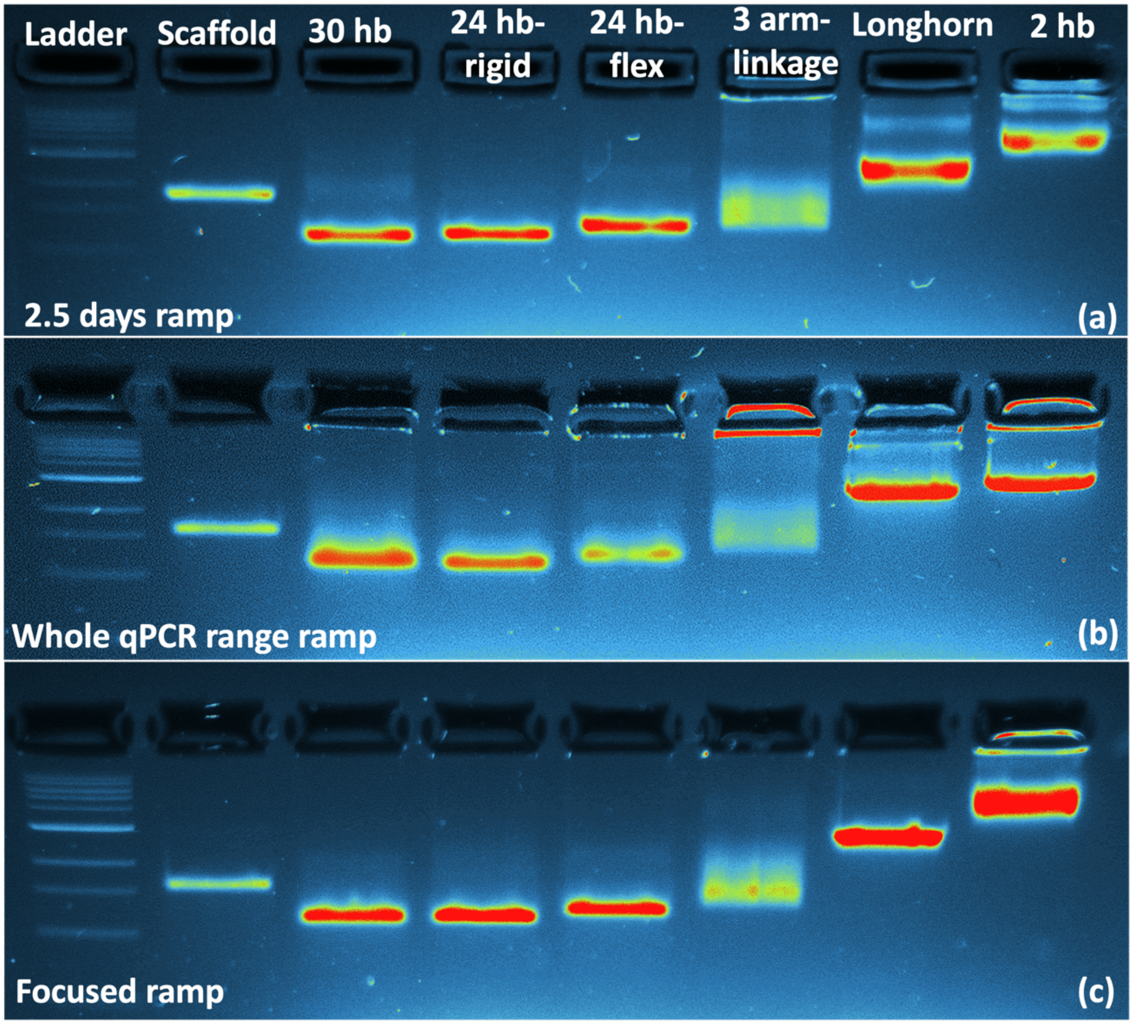


**Figure S6.** Agarose gel electrophoresis results in different structures after conducting a) 2.5 days thermal ramp, b) the real-time fluorometry ramp used in Figure 2 and c) focused thermal annealing at real-time fluorometry peaks regions and shock-freezing them after the last peak. The yield of the structures based on these gel images are presented in Figure 8.

(a)

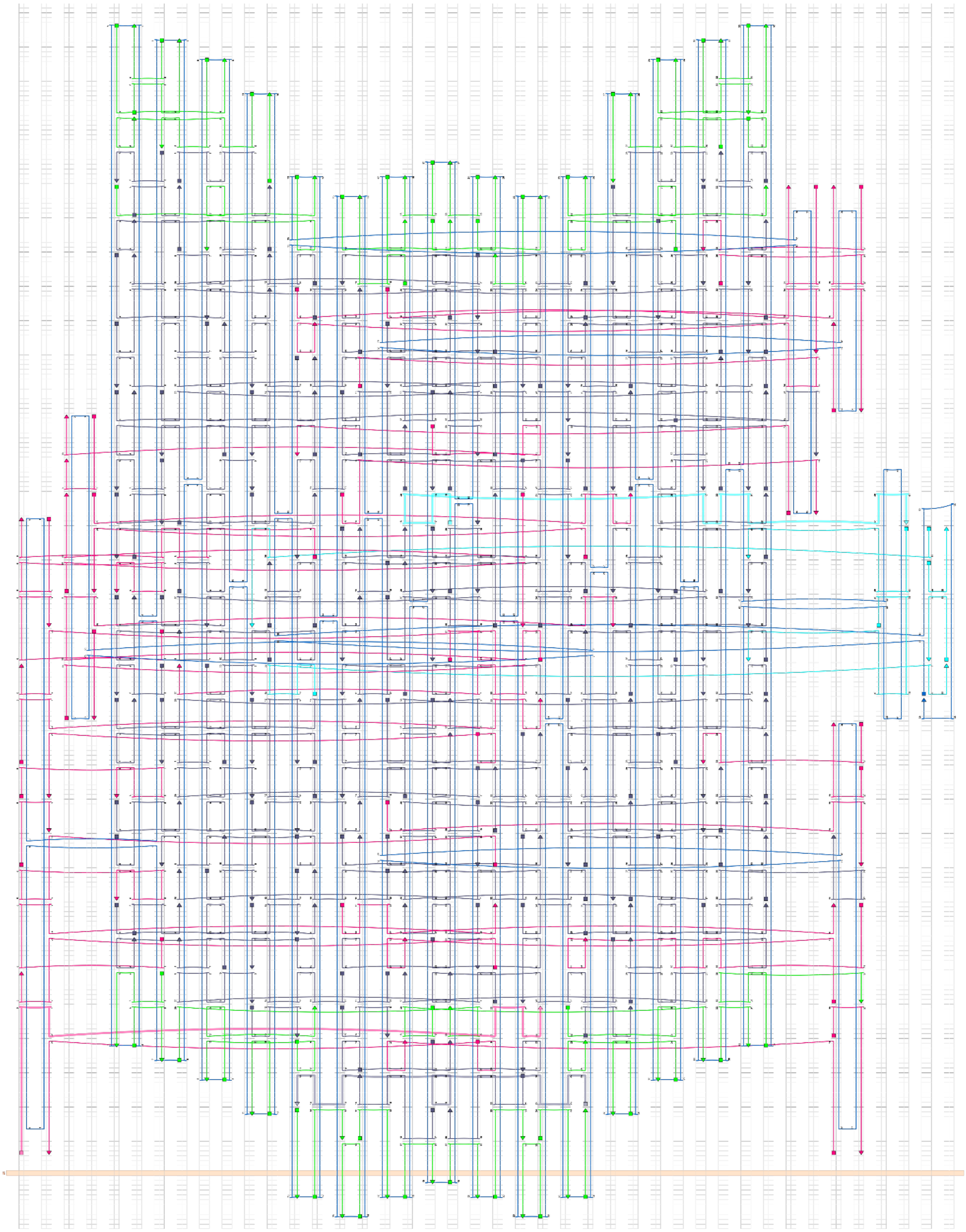

**(b)**

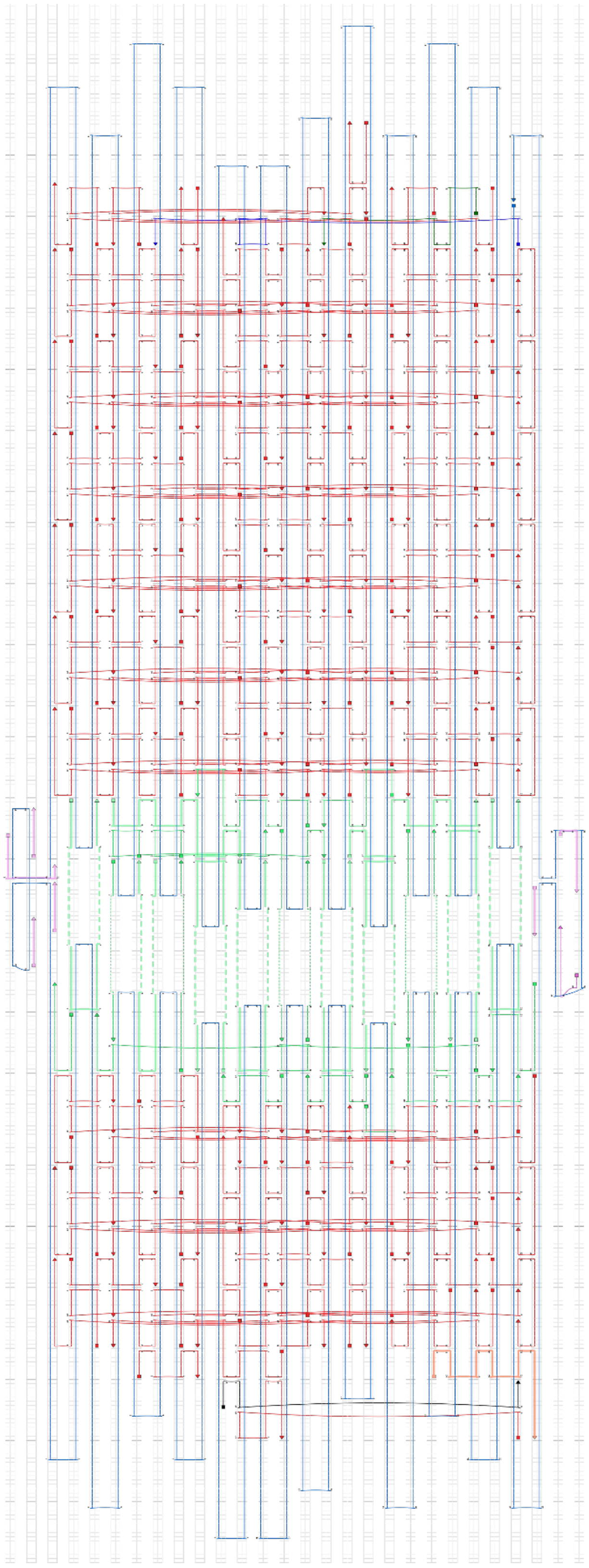

(c)

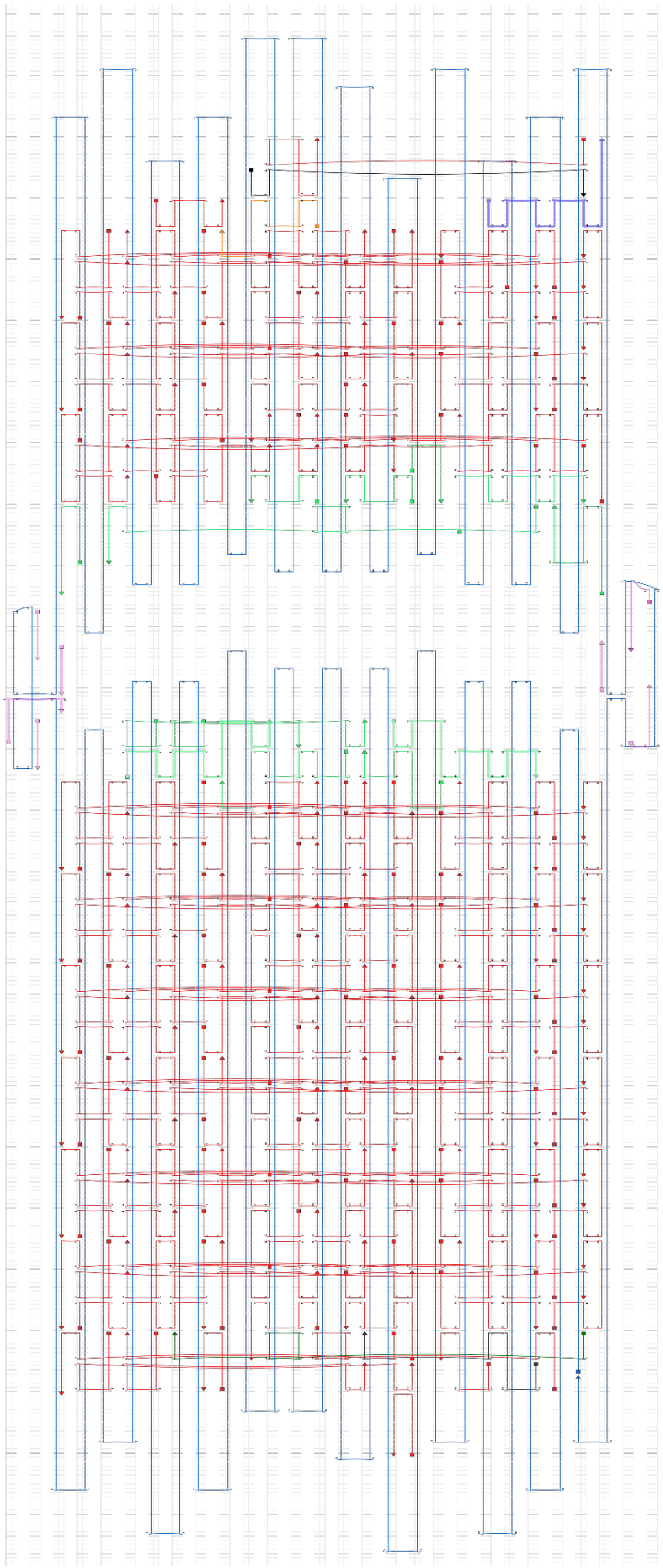

**(d)**

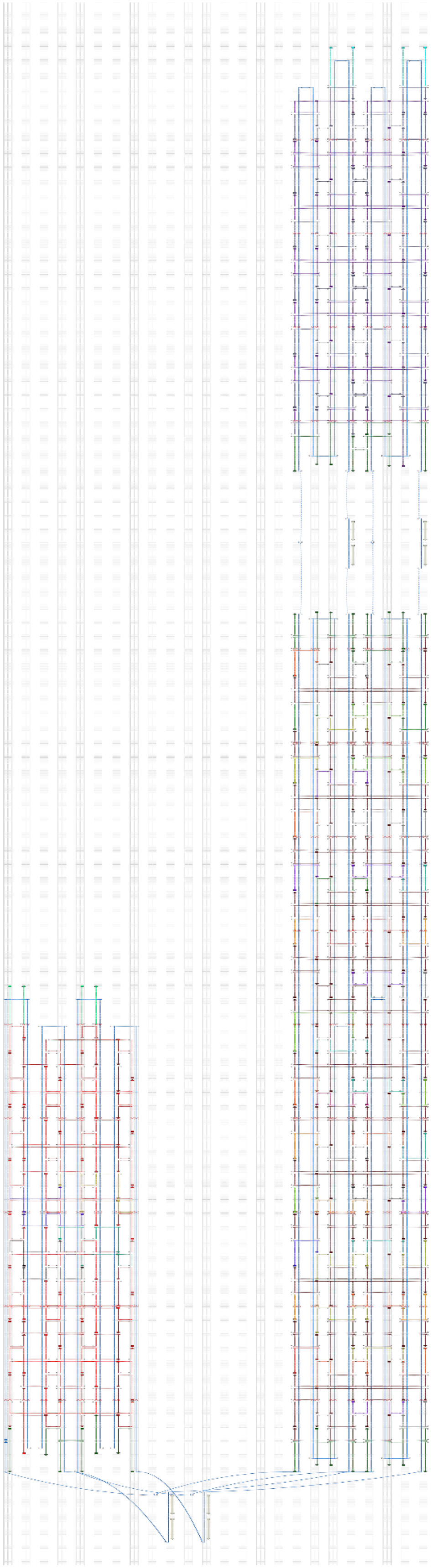

**(e)**

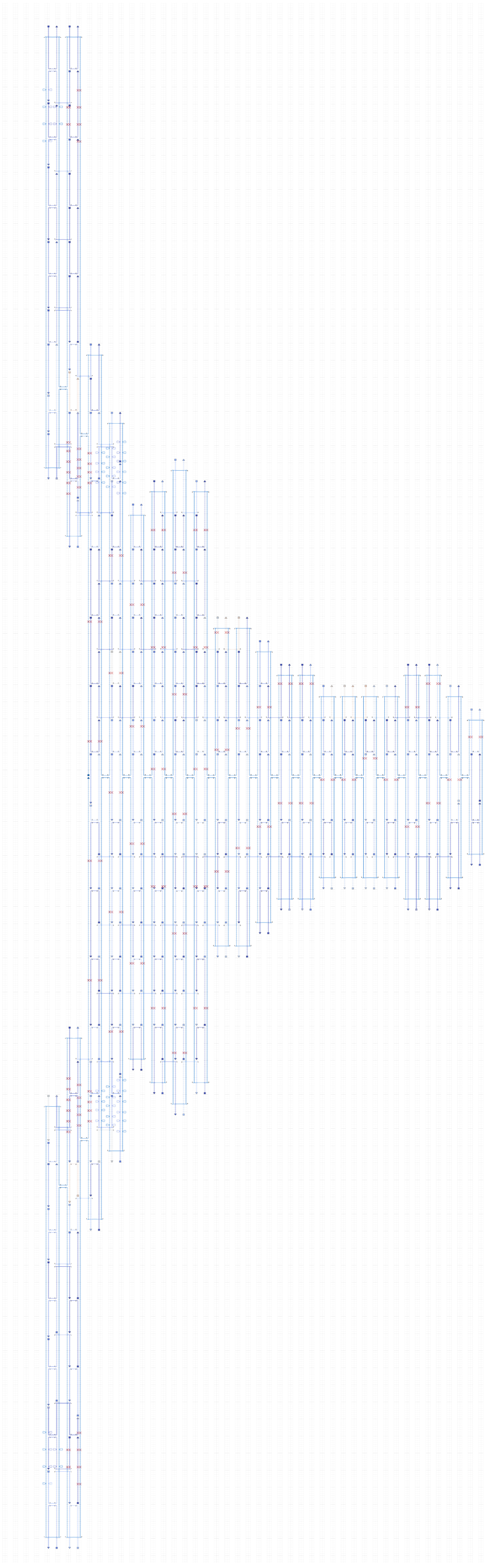

**(f)**

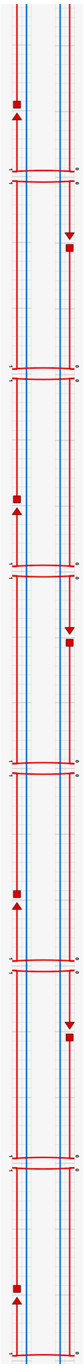

**Figure S7.** caDNAno designs for the studied DNA origami objects: (a) 30 hb, (b) 24 hb-rigid, (c) 24 hb-flex, (d) 3 arm-linkage, (e) Longhorn, and (f) cross-section of 2 hb.
